# Transcriptome and metabolome analysis provide insights into root and root released organic anion responses to phosphorus deficiency in oat

**DOI:** 10.1101/262014

**Authors:** Yanliang Wang, Erik Lysøe, Tegan Armarego-Marriott, Alexander Erban, Lisa Paruch, Andre van Eerde, Ralph Bock, Jihong Liu-Clarke

## Abstract

Root and root-released organic anions play important roles in uptake of phosphorus (P), an essential macronutrient for food production. Oat, ranking sixth in the world’s cereal production, contains valuable nutritional compounds and can withstand poor soil conditions. The aim of this research was to investigate root transcriptional and metabolic responses of oat grown under P-deficient and P-sufficient conditions. We conducted a hydroponic experiment and measured root morphology, organic anions exudation, and analysed changes in the transcriptome and metabolome, to understand oat root adaptation to P deficiency. We found that oat roots showed enhanced citrate and malate exudation after four weeks of P-deficiency. After 10 days of P-deficiency, we identified 9371 differentially expressed transcripts with a two-fold or greater change (*p* < 0.05): forty-eight sequences predicted to be involved in organic anion biosynthesis and efflux were consistently up-regulated; twenty-four up-regulated transcripts in oat were also found up-regulated upon P starvation in rice and wheat under similar conditions. Phosphorylated metabolites (i.e. glucose-6-phosphate, myo-inositol-phosphate) reduced dramatically, while citrate and malate, some sugars and amino acids increased slightly in P-deficient oat roots. Our data provide new insights into the root responses to P deficiency and root-released organic anions in oat.

**Highlight:** We found oat- a monocot food crop, showed high exudation rate of citrate under phosphorus deficiency; root transcriptome and metabolome were then investigated to understand oat adaptation to P deficiency.

## Introduction

Oat, *Avena sativa* L., is one of the most important food and feed crops in the world. It contains various nutritional and health-promoting compounds such as avenanthramides, tocopherols (vitamin E), and digestive fibre (β-glucans) (Gutierrez-Gonzalez *et al.*, 2013; Gutierrez-Gonzalez & Garvin, 2016), which may help reduce blood pressure and blood sugar levels, reduce cholesterol, and promote healthy gut bacteria (Alminger and Eklund-Jonsson, 2008; Nazare *et al.*, 2009; Valeur *et al.*, 2016; Whitehead *et al.*, 2014). In addition, oat can withstand poor soil conditions (e.g. acidic soils; Hill 1931; Stewart and McDougall, 2014) and is widely cultivated in temperate climates.

The world population is estimated to reach 9.8 billion by 2050 (www.un.org). In order to feed the world, we need to secure sustainable food production worldwide. Phosphorus (P) is a key macronutrient with significant impact on plant growth and productivity. The application of millions of tons of P fertilizers every year exhausts the limited and non-renewable P stocks available in the world (Cordell *et al.*, 2009). Hence, an understanding of the mechanisms of P mobilization and uptake in order to improve P acquisition efficiency, particularly in staple cereal food crops, is of importance for food security, and for environmentally friendly and sustainable food production in the future (Cordell *et al.*, 2009; Faucon *et al.*, 2015; Vance *et al.*, 2003).

Plants have evolved various adaptive strategies to cope with P deficiency in nature. Examples are morphological responses such as changes in root architecture (Hermans *et al.*, 2006; Lynch, 2011); physiological adaptations like secreted organic anions and acid phosphatases (Cheng *et al.*, 2014; Gahoonia *et al.*, 2000; Hedley *et al.*, 1982; Hoffland *et al.*, 1989; Jones 1998; Lambers *et al.*, 2006; Pang *et al.*, 2015; Ryan *et al.*, 2001; Wang *et al.*, 2016), biochemical responses to optimize utilization of internal P such as replacement of P-lipids with non-P lipids (Chiou & Lin, 2011; Faucon *et al.*, 2015; Lambers *et al.*, 2015; Plaxton & Tran, 2011;), and molecular responses like induced expression of high-affinity phosphate transporters (Wu *et al.*, 2013; Zhang *et al.*, 2014). In addition, for plant species that are capable of interacting with mycorrhizal fungi, P uptake by the mycorrhizal hyphae is the dominant pathway for P acquisition (Smith *et al.*, 2003; Sawers *et al.*, 2017).

Multiple genes and different mechanisms are required to improve plant tolerance to P deficiency. Thousands of plant genes that are differentially expressed in response to P deficiency have been identified by microarray, expressed sequence tags (EST) analysis and RNA sequencing (RNA-seq) in a large variety of plant species such as Arabidopsis (*Arabidopsis thaliana*), potato (*Solanum tuberosum*), rice (*Oryza sativa*), wheat (*Triticum aestivum*), and white lupin (*Lupinus albus*) (Hammond *et al.*, 2011; Misson *et al.*, 2005; Oono *et al.*, 2011; O’Rourke *et al.*, 2013; Oono *et al.*, 2013). Regulatory components identified include transcription factors (TFs such as PHR1), SPX domain-containing proteins, plant hormones, microRNAs, protein modifiers and epigenetic modifications (Chiou & Lin, 2011; Lin *et al.*, 2009; Panigrahy *et al.*, 2009; Wu *et al.*, 2013; Yang & Finnegan, 2010; Zhang *et al.*, 2014). The networks of regulatory genes that are necessary to sense and respond to P deficiency are complex and differ in different plant species. For Poaceae species, the molecular mechanisms associated with P uptake, translocation and remobilization are well elucidated in rice (Panigrahy *et al.*, 2009; Oono *et al.*, 2013; Wu *et al.*, 2013). Briefly, P starvation activates expression of OsPHR2, which triggers gene expression of phosphate transporters, purple acid phosphatase (PAP) and other proteins contributing to enhanced P uptake. In addition, SPX domain-containing proteins, which are activated by expression of OsPHR2, support maintenance and utilization of internal phosphate. OsPHO1 functions in P translocation (xylem loading) while SIZ1 (a small ubiquitin like modifier SUMO E3 ligase) targets PHR2 and acts both negatively and positively on various P deficiency responses. Finally, microRNA399 targets PHO2 to regulate plant P homeostasis (Chiou and Lin, 2011; Oono *et al.*, 2013; Panigrahy *et al.*, 2009; Wu *et al.*, 2013).

Plant root plays an essential role in P uptake. It is well known that to promote P uptake at reduced P availability, most species allocate more biomass to roots, increase root length and develop more and longer root hairs and later roots and so on (Hermans *et al.*, 2006; Lambers *et al.*, 2006; Lambers *et al.*, 2015; Lynch, 2011). Accordingly, mechanisms regulating root architecture like phytohormones and particularly the auxin responses under P limitation have been elucidated, as well as genes associated with those responses (see reviews Chiou and Lin, 2011; Lin *et al.*, 2009; Lynch, 2011; Panigraphy *et al.*, 2009). Root exuded organic anions is also considered as an important mechanism to mobilize soil less-available P and enhance plant P uptake, while there is little information on the molecular mechanisms involved in organic acids biosynthesis and efflux under P deficiency.

Despite the importance of oats, limited research has been carried out on its adaptation to P starvation, and particularly the molecular regulation of root and root-released organic anions in response to P deficiency. In a previous report, we found that oat showed an increased root mass / total biomass ratio, high percentage of root colonization by arbuscular mycorrhizal fungi (AMF), large amounts of rhizosphere organic anions and efficient P uptake in low P availability soils (Wang *et al.*, 2016). These findings paved the way for our current study on the molecular mechanisms underlying P deficiency responses in oat roots and the genes and metabolites involved. Here, we compared gene expression and metabolome profiles of oat roots exposed to P sufficiency (100 μM KH_2_PO_4_) and deficiency (1 μM KH_2_PO_4_) conditions by hydroponic culture. The objectives were: 1) identification of differentially expressed transcripts in oat roots in response to P deficiency, with particular focus on up-regulated transcripts associated with organic anions biosynthesis and exudation; 2) discovery of conserved responsive genes in rice, wheat and oat, and transcripts unique for oat; 3) assessment of differential metabolite accumulation in response to P deficiency and the transcriptional program triggered by it. The overall goal of the current study was to identify candidate genes that will enrich our understanding of oat adaptation to P deficiency and that may be useful to future breeding and genetic engineering efforts towards oat improvement.

## Materials and Methods

### Plant growth and harvest

Seeds of the oat cultivar ‘BELINDA’ were germinated and grown hydroponically in full strength nutrient media (Wang *et al.*, 2015) in the greenhouse at the Norwegian University of Life Sciences. Fourteen days after sowing, seedlings were transferred to the same medium supplemented with 100 (P100) or 1 (P1) μM KH_2_PO_4_, respectively, and pH adjusted to 5.8±0.2. Plants were grown under a photoperiod of 16 h light and 8 h dark at a light intensity of 200 ± 20 μmol m^−2^ s^−1^ and 50-75% relative humidity, with a temperature of 25 ˚C/ 16 ˚C (day/ night). The nutrient solution was replaced every third day.

Ten days post-treatment, four root samples (representing four independent biological replications) from both P100 and P1 treated plants were collected for RNA extraction and analysis as described by Oono *et al.* (2011). The sampled roots were mixed samples containing the root cap zone, elongation zone and a part of the maturation zone. Those eight root samples, together with another eight samples (four from P1 and four from P100) were used for root metabolome analysis. When sampled roots for RNA and metabolite extraction, the roots were quickly washed and water-removed and then immediately placed in liquid nitrogen. Additionally, eight plants (two treatments, four replicates) were used for studies of root morphology, root organic anions and biomass determinations after four weeks of different P treatments. All the plants were in vegetative growth phase, with tiller but before heading.

### Root released organic anions, root morphology and biomass determination

For root exudate collection, briefly, whole root systems of intact plants were carefully washed with deionized water to remove the nutrient solution. The whole root system was then placed into ultrapure Milli-Q water (Millipore, Billerica, MA, USA) in a container to collect root exudates (Khorassani *et al.*, 2011; Wang *et al.*, 2015). Afterwards, micropur (0.01 g L^−1^, Katadyn Products, Kemptthal, Switzerland) was added to the solution to inhibit the activity of microorganisms (Cheng *et al.*, 2014). The collected root exudates were analysed by liquid chromatography triple quadrupole mass spectrometry (LC–MS/MS), as described in a previous study (Wang *et al.* 2015). Root released organic anions were collected and analysed after plants had been grown hydroponically under P1 or P100 for 2 weeks and 4 weeks, respectively.

After 4 weeks, the total number of green leaves and senesced leaves was recorded. For root morphology determination, WinRHIZO (EPSON 1680, WinRHIZO Pro2003b, Regent Instruments Inc., Quebec, Canada) was used to measure root length, number of lateral roots and root surface area. Shoot and root dry weight (DW) were measured separately after being oven-dried for 48 h at 65 °C. Shoot P concentrations were subsequently determined by inductively coupled plasma atomic emission spectroscopy (Wang *et al.*, 2015).

### RNA extraction and quality control

Total RNA was extracted using a Spectrum™ Plant Total RNA Kit (Sigma-Aldrich, St. Louis, MO, USA) and genomic DNA was removed using On-column DNase I digest kit (Sigma-Aldrich, St. Louis, MO, USA). RNA quantity and quality was assessed by a NanoDrop spectrophotometer (NanoDrop Technologies, Wilmington, DE, USA) and Agilent 2100 Bioanalyzer (Santa Clara, CA, USA). RNA samples with RIN (RNA integrity number) scores greater than 9.0 were used for RNA-Seq. Eight independent root cDNA libraries were prepared according to Illumina’s TruSeq® RNA Sample Preparation v2 Guide and 125-bp paired-end reads were sequenced using an Illumina HiSeq 2500 sequencer (Illumina, San Diego, CA, USA) at the Norwegian Sequencing Centre (www.sequencing.uio.no).

### Sequence processing and analysis

A total of 215,087,481 paired-end short read sequences were quality checked, trimmed and *de novo* assembled using CLC Genomics Workbench v9.01 (QIAGEN Aarhus, Denmark), generating 207017 contigs, with a maximum contig length of 13,319 nt, a minimum contig length of 200 nt and a mean contig length of 801 nt (Table 1). Gene expression was calculated and normalized using RPKM (Reads Per kb per Million reads). Differential expression between P1 and P100 was analyzed by *t*-tests and up-/down- regulation of genes was considered to be significant if equal or greater than two-fold (*p* < 0.05). Totally, 41,679 transcripts were filtered out and selected to be *de novo* Oat Root Transcriptome (*dn*ORT), used as a reference for further analysis. The *dn*ORT sequences were a combination of the differentially expressed transcripts, and other transcripts with RPKM ≥ 1.5 regardless of P treatments. These sequences were annotated using Blast2Go (Conesa *et al.*, 2005) and MapMan (Thimm *et al.*, 2004). RNA sequencing raw data were deposited to the GeneBank Sequence Read Archive (SRA) database under bioproject identifier PRJNA3 55647.

**Table 1.** Transcriptome statistics.

|  |  |
| --- | --- |
| Total number of reads | 215,087,481 |
| Assembled contigs | 207,017 |
| Minimum length (nt) | 200 |
| Maximum length (nt) | 13,319 |
| Mean length (nt) | 801 |
| Contigs larger than 1,000 nt | 9631 |
| Up-regulated contigs | 7,817 |
| Down-regulated contigs | 1,554 |
| Annotated contigs | 41,679 |

nt: nucleotides

### Real-time quantitative reverse transcription PCR (qRT-PCR) analysis

First-strand cDNA was synthesized from 1.0 µg RNA using iScript^TM^ Adv cDNA kit for qRT-PCR (Bio-Rad, USA). The qRT-PCR reactions were carried out on CFX96 Real-time system (Bio-Rad, USA) using SsoAdvanced^TM^ Universal SYBR^®^ Green Supermix (Bio-Rad, USA) with transcript-specific primers shown in Supporting Information Table S5. Ten ng cDNA were used as template in a 20 µl qPCR reaction consisting of 0.8 µM primers. After initial denaturing at 95°C for 5 min, the reaction was followed by 40 cycles at 95°C for 15 s, 61 °C for 15 s and 72°C for 45 s. The expression of endogenous reference genes *EF1α* (Elongation factor 1*α*, Kemen *et al*, 2014) and *β-Actin* was used to normalize the expression level estimated by the ΔΔCq method provided by CFX Manager 3.1 (Bio-Rad, USA). Four biological replicates of each treatment and three technical replicates of each sample were applied in the analysis. The qPCR data were represented as fold change (P1 mean value: P100 mean value) derived from relative normalized expression level from four biological replicates and further compared with RNA-seq results (P1 RPKM means/ P100 RPKM means). *R* software (version 3.2.2) and one-way ANOVA were used to examine significant differences between P1 and P100 treatments. Heat maps were generated in Heml 1.0: Heatmap illustrator as described by Deng *et al.* (2014).

### Root metabolite extraction

Eight replicate samples each of roots from plants grown under P1 and P100 conditions were sampled. Frozen samples (with water content varying between 95.3% and 96.6%) of 100 mg (± 10%) root were ground to homogeneity (2 min, 30 Hz; Grinding Mill MM310; Retsch, DE) under frozen conditions. To each sample was added 360 μL precooled (-20 ºC) extraction buffer (300μL methanol, 30 μL 2 mg mL^−1^ nonadecanoic acid methylester in chloroform, 30 μL 0.2 mg mL^−1^ 13-C sorbitol in methanol) and samples shaken for 15 min, 70 ºC, 1000 rpm (Thermomixer Comfort Eppendorf, DE). Samples were cooled to room temperature, added to 200 μL CHCl_3_ and further shaken for 5 min, 37 ºC, 1000 rpm. To each sample was added 400 μL H_2_O and, following vortexing, samples were centrifuged (5 min, 14000 rpm) to facilitate phase separation. Finally, 160 μL of the upper, polar phase was aliquoted and dried overnight by Speed Vac.

### Gas chromatography–mass spectrometry (GC-MS) metabolite profiling and identification

Prior to gas chromatography–electron impact–time of flight mass spectrometry (GC-EI/TOF-MS) analysis, metabolites were methoxyaminated and trimethylsilylated. Briefly, addition of 40 µl MeOX (40 mg ml^−1^ methoxyaminhydrochloride in pyridine), samples were shaken (1.5 h, 30 ºC), addition of 80 µl BSTFA mixture (70 µl N, O-Bis(trimethylsilyl)trifluoroacetamide +10 µl Alkane-Mix (n-alkanes: C10, C12, C15, C18, C19, C22, C28, C32, and C36)), and further shaken (30 min, 37 ºC). GC-MS was undertaken using an Aligent CP9013 column in an Agilent 6890N24 gas chromatograph, coupled to a Pegasus III, similar to that previously described (Dethloff *et al.*, 2014; Erban *et al.*, 2007; Wagner *et al.*, 2003). Measurements were undertaken both splitless and split (1/30) from an injection volume of 1 µL, with bulk metabolites reaching the upper detection limit in split-less measurements evaluated from split data. Retention indices were calibrated based on added n-alkanes (Strehmel *et al.*, 2008).

Chromatograms were visually controlled, baseline corrected and exported in NetCDF format using ChromaTOF. Further data processing and compound identification was performed with TagFinder (Luedemann *et al.*, 2008) and by matching to the Golm Metabolome Database (GMD, http://gmd.mpimpgolm.mpg.de/; Kopka *et al.*, 2005; Schauer *et al.*, 2005), and the NIST08 database (http://www.nist.gov/srd/mslist.htm). Manually supervised metabolite annotation and quantification was undertaken with the requirement of at least three specific quantitative mass fragments per compound, and a retention index deviation < 1.0%. Data were normalized to sample fresh weight, the internal 13-C6 sorbitol, and represented relative to the mean value of P100 samples per analyte.

### Principal component analysis (PCA) and statistical analysis

PCA was carried out by using the program *R*. Data were normalized to the median of the P100 samples and subsequently subjected to logarithmic (Log2) transformation. Missing values were not substituted with zero or a constant value. Statistical testing was performed using the *t* test in multiple experiment viewer MeV (Saeed *et al.*, 2006), based on Log2 transformed data, followed by Mann-Whitney false discovery rate (FDR) correction at *α*< 0.05, due to non-normal distribution (as shown by PCA analysis) of the metabolite data.

## Results

### Plant growth, root morphology and root released organic anion analysis

The plant phenotype and root released organic anions were examined to study the effects of P deficiency on oat growth and root exudates under hydroponic condition. After four weeks of growth under two different P regimes (1 and 100 μM KH_2_PO_4_), a drastic reduction of shoots P concentration was observed in plants grown under P deficient conditions (P1), compared with P100 (0.90 vs 6.35 mg g^−1^). Oat plants subjected to P1 treatment showed on average a 55% reduction in the total number of leaves, 44% more senescent leaves (number of senescent leaves: number of total leaves), 68% less shoot dry biomass and 96% greater root mass ratio (root dry biomass: total dry biomass) than P100 plants as shown in Fig. 1A-D. Moreover, P1 treatment plants showed shorter total root length (25%), less root surface area (27%) and more lateral roots (14%) than P100 plants (Fig. 2A-D). Furthermore, after four weeks, compared with P100 treated plants, which showed no detectable root released organic anions, P1 roots had higher exudation rates of citrate and malate, 927 and 81 nmol h^−1^ g^−1^ root dry weight (DW) respectively, as shown in Fig. 2E. By contrast, no organic anions were detected in either P1 or P100 root exudates collected after two weeks treatment.

**Figure 1.**
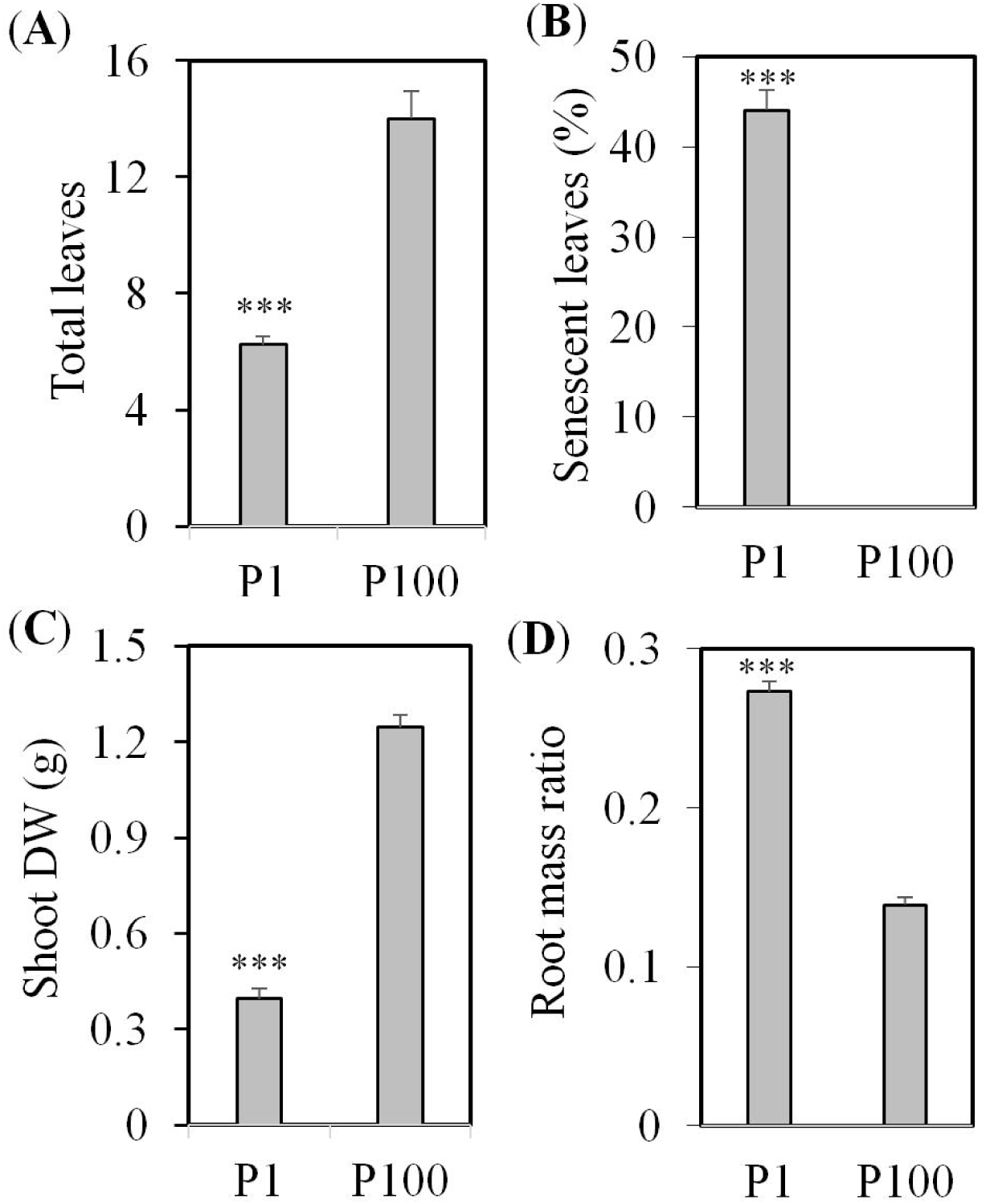
Plant growth response to P1 and P100 treatments. (**A**) Total leaves, (**B**) Senescent leaves, (**C**) Shoot dry weight, and (**D**) Root mass ratio. Error bars indicate SE (n = 4). Significant differences are indicated (ns, not significant; ***, *p* < 0.001).

**Figure 2.**
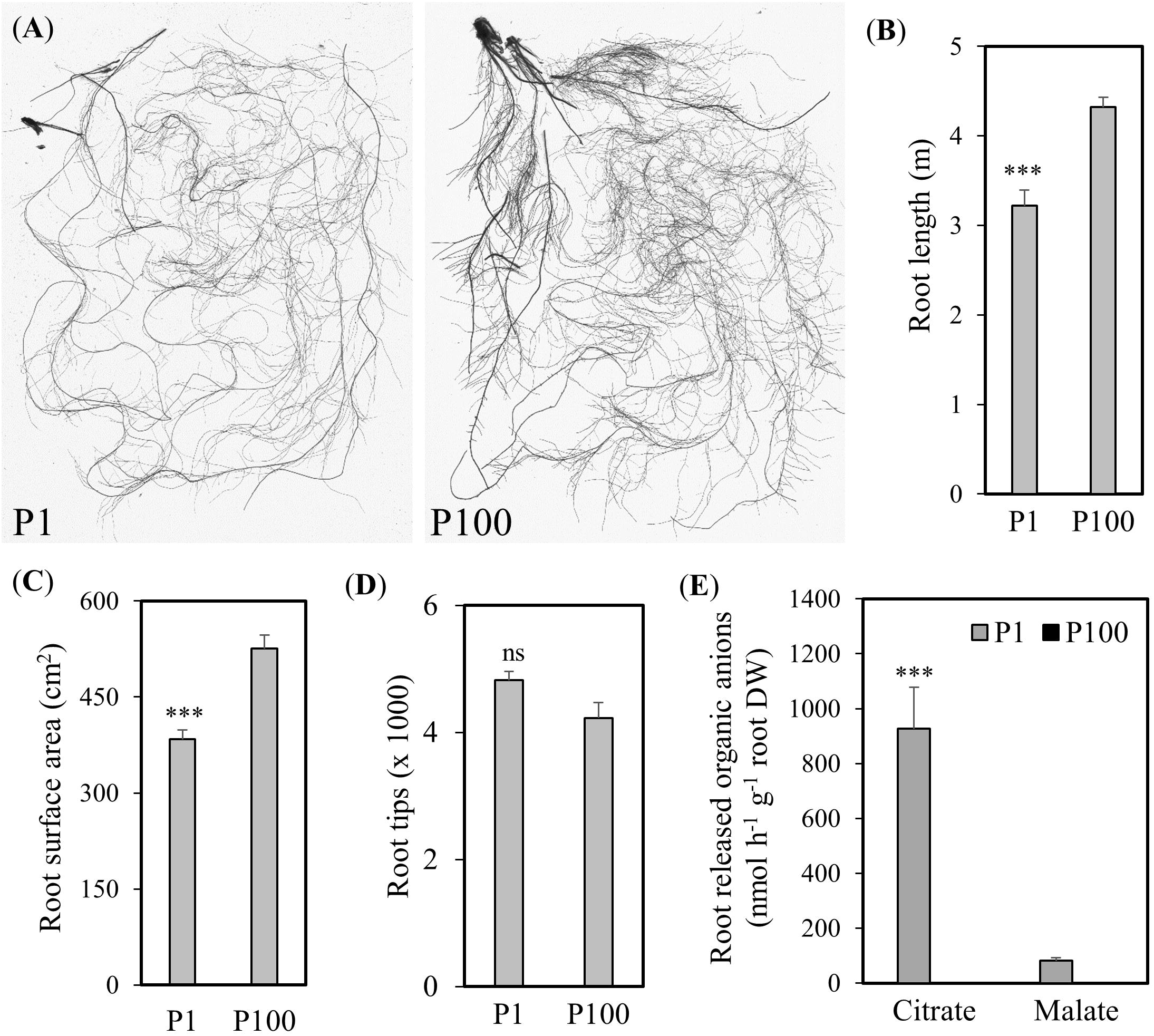
Oat root response to P1 and P100 treatments. (**A**) Representative photos of root structure, (**B**) Root length, (**C**) Root surface area, (**D**) Number of root tips, and (**E**) Root released organic anions. Error bars indicate SE (n = 4). Significant differences are indicated (ns, not significant; ***, *p* < 0.001).

### Transcriptome analysis of root response to P deficiency

Next, RNA-seq was performed to evaluate gene expression from P deficient roots of oat plants as compared to P sufficient roots. We did Blastx and GO analyses after the *dnORT* database was constructed as described in the Material and Methods part. Approximately 9.4% of the *dn*ORT transcripts could not be assigned to any Blastx hits (E-value > 1E-3) as shown in Supplementary Data Fig. S1. The blastx top hit species were: *Brachypodium distachyon* (22%), *Hordeum vulgare* subsp. *vulgare* (17%), *Aegilops tauschii* (15%), *Triticum urartu* (10%) and *Oryza sativa japonica* group (4%) (Fig. S2). Functional gene ontology (GO) classification of *dn*ORT sequences suggested that the biological process was mainly represented by the term ‘cellular and metabolic processes’, and the most represented GO subcategories within the cellular component main term were ‘cell or cell part’ and ‘membrane or membrane part’. When the sequences were categorized according to the molecular function main term, 10,699 transcripts corresponded to ‘binding category’ and 10,522 sequences to ‘catalytic activity’ (Fig. S3). Putative functions (with InterProScan) were predicted for 62% of the sequences (Fig. S4).

In total, 9,371 transcripts (7,817 up-regulated, 1,554 down-regulated) were differentially expressed in response to P deficiency as shown in Table 1. Gene ontology (GO) categories showed that up-regulated transcripts under P deficiency were categorized into more than 40 groups, such as oxidation-reduction process, transmembrane transport, carbohydrate metabolic process, response to osmotic stress, biosynthetic process, pyruvate metabolism, tricarboxylic acid cycle, acid phosphatase, and CCAAT-binding complex (Fig. 3).

**Figure 3.**
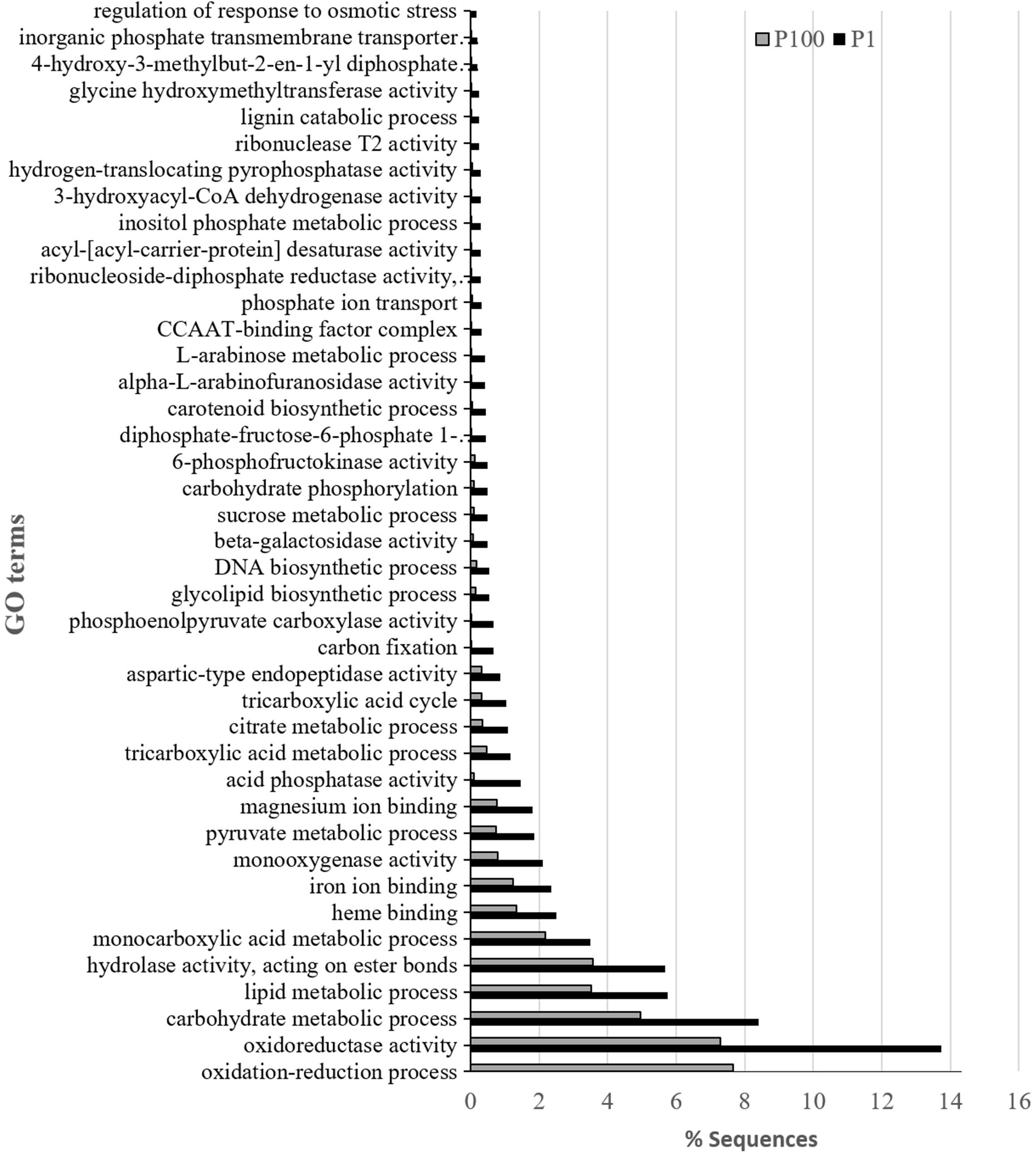
Functional annotation of up-regulated sequences based on gene ontology (GO) categorization. *y*-axis indicates the category, *x*-axis the percentage of transcripts in a category.

Reciprocal tblastx (E < 1E-10) analysis showed that 24 oat transcripts which were up-regulated in P1 matched the conserved responsive genes previously found up-regulated in both rice and wheat (Oono *et al.*, 2013) as listed in Table 2. We also found 25 unique responsive transcripts (between 308 nt and 1672 nt) in oat roots that were up-regulated more than two-fold (*p* < 0.001, P1 RPKM means > 5), without any blast hits in currently available databases, i.e. they seem to be exclusively expressed in oat (Table 3).

**Table 2.** Conserved responsive transcripts found in oat, wheat and rice under P deficiency.

| *dn*ORT ID | Rice proteins | Wheat_proteins (TRIAE_CS42) | P1<br>RPKM | P100<br>RPKM | Fold change<br>P1/P100 | Gene | Gene annotation |
| --- | --- | --- | --- | --- | --- | --- | --- |
| Contig000022 | Os05t0137400-01 | 1AS_TGACv1_020885_AA0080310.2.1 | 107.9 | 48.1 | 2.2 |  | Similar to aspartic protease precursor |
| Contig000835 | Os05t0387200-01 | 1DL_TGACv1_061485_AA0196630.2.1 | 13.5 | 1.7 | 8.0 | SQD1 | Sulphite: UDP-glucose sulfotransferase |
| Contig010807 | Os10t0500600-01 | 1DL_TGACv1_062297_AA0212030.5.1 | 48.3 | 22.6 | 2.1 |  | Zinc finger, C2H2-like domain containing protein |
| Contig011660 | Os07t0100300-02 | 2AL_TGACv1_094153_AA0293430.1.1 | 17.4 | 1.3 | 13.4 |  | Glycosyl transferase, group 1 domain containing protein |
| Contig009578 | Os10t0100500-01 | 2AS_TGACv1_113238_AA0353330.1.1 | 61.3 | 11.4 | 5.4 |  | Serine/threonine protein kinase-related domain containing protein |
| Contig001778 | Os07t0622200-01 | 2AS_TGACv1_113290_AA0354140.2.1 | 71.8 | 68.0 | 1.1 |  | Similar to M-160-u1_1 |
| Contig009737 | Os07t0630400-01 | 2BS_TGACv1_146583_AA0468610.1.1 | 5.2 | 1.1 | 4.8 | OsRNS1 | Ribonuclease T2 family protein |
| Contig005229 | Os04t0555300-01 | 2DL_TGACv1_158105_AA0509380.4.1 | 41.6 | 9.1 | 4.6 |  | Similar to glycerol 3-phosphate permease |
| Contig019750 | Os04t0652700-01 | 2DL_TGACv1_158583_AA0522480.2.1 | 4.8 | 1.2 | 4.0 |  | Similar to nuclease PA3 |
| Contig061747 | Os01t0128200-01 | 3AS_TGACv1_210696_AA0677330.2.1 | 4.0 | 1.4 | 2.8 |  | Similar to nuclease I |
| Contig013859 | Os01t0897200-04 | 3B_TGACv1_224141_AA0792910.2.1 | 68.9 | 21.8 | 3.2 | OsRNS2 | Ribonuclease 2 precursor |
| Contig004306 | Os06t0115600-01 | 4AL_TGACv1_289135_AA0965550.1.1 | 76.7 | 53.6 | 1.4 |  | Similar to CYCLOPS |
| Contig026884 | Os08t0299400-01 | 4AL_TGACv1_289998_AA0980080.1.1 | 13.9 | 0.0 |  | MGD | MGDG synthase type A |
| Contig003298 | Os03t0238600-01 | 4AS_TGACv1_308481_AA1028160.1.1 | 273.2 | 54.6 | 5.0 | PAP | Similar to purple APase |
| Contig007532 | Os09t0553200-01 | 5AL_TGACv1_374888_AA1211020.2.1 | 504.8 | 181.3 | 2.8 | UGPase | UDP-glucose pyrophosphorylase |
| Contig003667 | Os09t0478300-01 | 5AL_TGACv1_376126_AA1232370.2.1 | 17.7 | 7.5 | 2.4 |  | Conserved hypothetical protein |
| Contig026720 | Os12t0554500-00 | 5AS_TGACv1_393365_AA1271860.2.1 | 11.6 | 0.1 | 167.7 |  | Lipase, class 3 family protein |
| Contig116061 | Os09t0379900-02 | 5BL_TGACv1_404442_AA1299920.1.1 | 1.6 | 0.6 | 2.5 |  | Endo-1,3(4)-beta-glucanase 2 like |
| Contig073770 | Os08t0433200-01 | 5BL_TGACv1_404654_AA1307490.1.1 | 28.0 | 5.4 | 5.1 |  | Conserved hypothetical protein |
| Contig007245 | Os09t0315700-01 | 5BL_TGACv1_407230_AA1354660.1.1 | 58.1 | 20.4 | 2.9 |  | Phosphoenolpyruvate carboxylase family protein |
| Contig000670 | Os02t0809800-01 | 6BL_TGACv1_501820_AA1620890.2.1 | 70.5 | 22.9 | 3.1 | PHO1:H2 | Root-to-shoot inorganic phosphate (Pi) transfer |
| Contig000259 | Os06t0178900-01 | 7BS_TGACv1_592527_AA1939830.5.1 | 184.6 | 82.2 | 2.2 |  | Vacuolar H<sup>+</sup> -pyrophosphatase |
| Contig027611 | Os05t0489900-01 | U_TGACv1_641100_AA2085080.2.1 | 13.9 | 6.4 | 2.2 |  | Calcium/calmodulin-dependent protein kinase |
| Contig044047 | Os09t0321200-00 | U_TGACv1_642666_AA2121200.1.1 | 0.9 | 0.1 | 11.0 |  | Similar to carotenoid cleavage dioxygenase |

**Table 3.** Unique P responsive transcripts found in oat.

| Oat transcript ID | *p*-value | P1 - RPKM | P100 - RPKM | Fold change<br>P1/P100 | Transcript length<br>(nt) |
| --- | --- | --- | --- | --- | --- |
| Contig000170 | 0.0001 | 339.8 | 7.6 | 44.5 | 486 |
| Contig001596 | 0.0005 | 27.8 | 12.0 | 2.3 | 582 |
| Contig002497 | 0.0001 | 10.3 | 1.4 | 7.5 | 452 |
| Contig005093 | 0.0007 | 193.8 | 42.9 | 4.5 | 567 |
| Contig006100 | 0.0001 | 80.8 | 2.0 | 39.6 | 308 |
| Contig006686 | 0.0000 | 60.8 | 1.0 | 59.6 | 400 |
| Contig007919 | 0.0002 | 67.4 | 0.5 | 134.2 | 586 |
| Contig009266 | 0.0003 | 5.2 | 0.1 | 35.3 | 1017 |
| Contig010063 | 0.0001 | 32.0 | 13.9 | 2.3 | 1256 |
| Contig011518 | 0.0000 | 16.3 | 0.6 | 28.0 | 978 |
| Contig022489 | 0.0000 | 13.6 | 0.0 | 907.5 | 1198 |
| Contig023173 | 0.0005 | 116.7 | 47.5 | 2.5 | 410 |
| Contig024878 | 0.0002 | 52.3 | 23.6 | 2.2 | 454 |
| Contig024909 | 0.0005 | 7.3 | 0.1 | 81.8 | 1004 |
| Contig029950 | 0.0001 | 8.9 | 0.4 | 24.7 | 817 |
| Contig034195 | 0.0003 | 17.7 | 8.4 | 2.1 | 812 |
| Contig035653 | 0.0004 | 5.5 | 1.7 | 3.1 | 702 |
| Contig039702 | 0.0005 | 8.3 | 0.7 | 11.6 | 447 |
| Contig040117 | 0.0000 | 14.3 | 0.2 | 77.4 | 630 |
| Contig040132 | 0.0000 | 7.2 | 0.0 | -- | 402 |
| Contig045423 | 0.0000 | 7.0 | 0.0 | -- | 410 |
| Contig045562 | 0.0005 | 5.7 | 1.5 | 3.7 | 1672 |
| Contig052573 | 0.0001 | 5.4 | 2.2 | 2.4 | 673 |
| Contig068492 | 0.0001 | 23.4 | 3.2 | 7.3 | 474 |
| Contig068533 | 0.0001 | 6.4 | 2.4 | 2.7 | 1167 |

Furthermore, as shown in Fig. 4 and Tables S1, S2, among the 7817 up-regulated transcripts, 128 transcripts were annotated as transcription factors (TFs), 57 sequences assigned as acid phosphatases and 18 as phosphate transporters. In addition, there were two sequences similar to SIZ1, SPX domain-containing proteins and PHO1 (which transfers P from roots to shoots), respectively, and one sequence was annotated as PHO2. Transcripts associated with auxin responses that regulate root development, and with disease and fungus responses, were also detected, as shown in Supporting Information Table S3.

**Figure 4.**
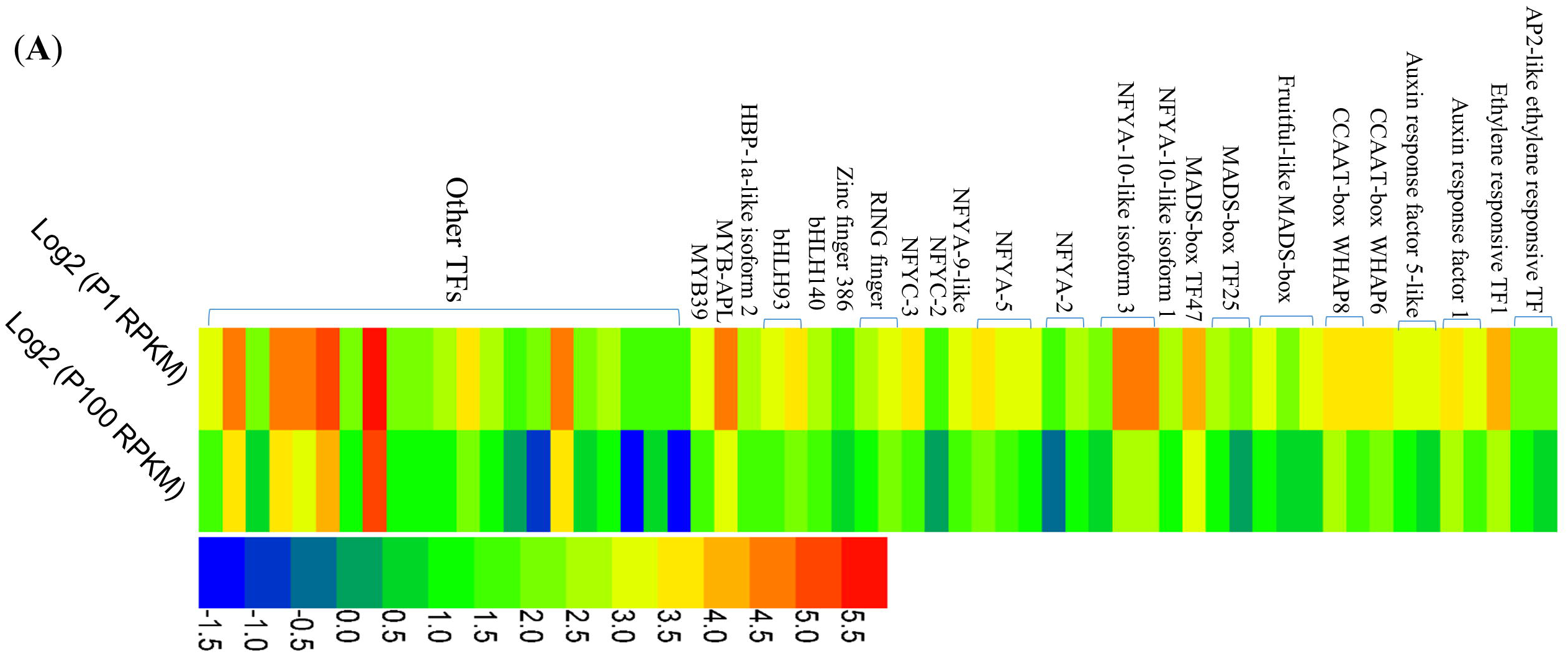
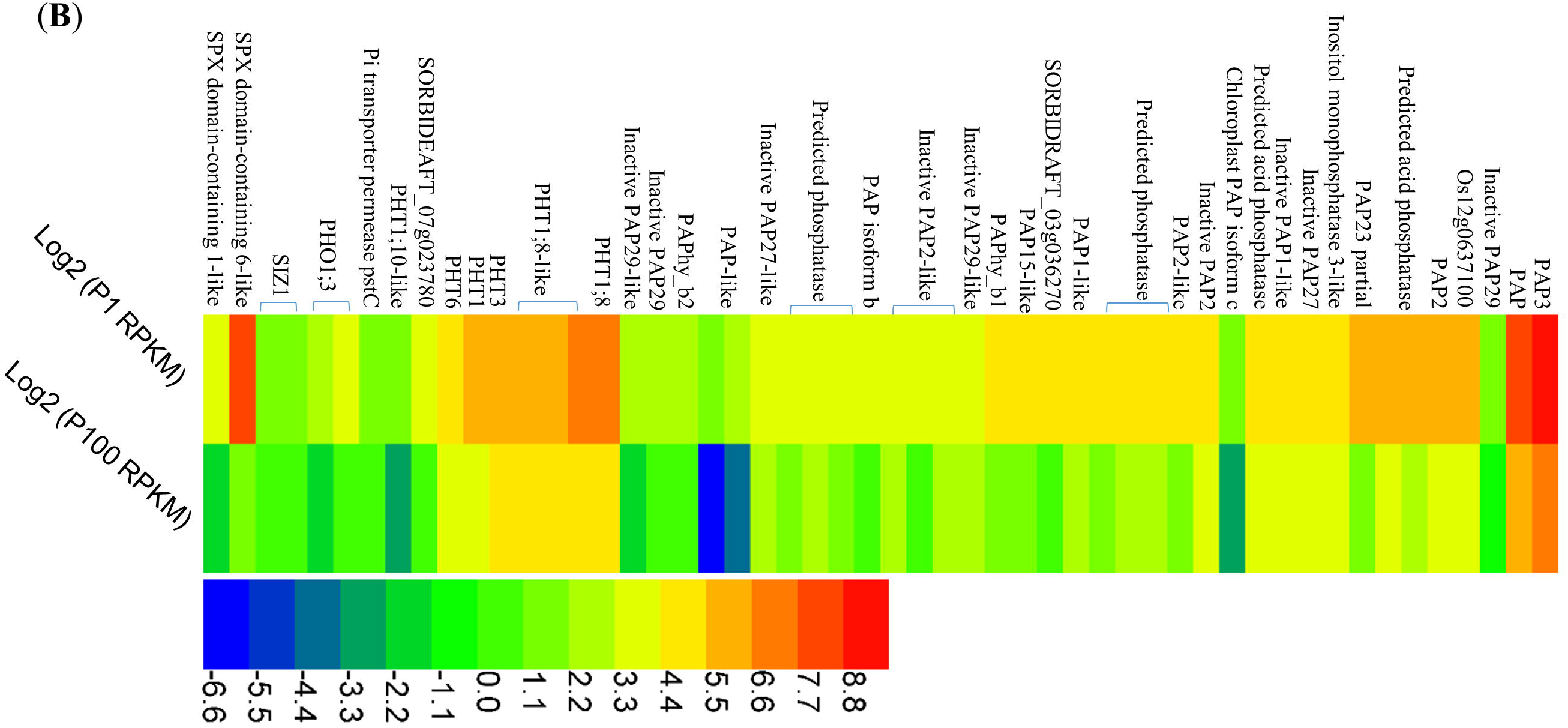
Heat map of expression profiling of up-regulated transcription factors (TFs) and selected known genes related to P deficiency. P1 induced up-regulated (*p* < 0.05) TFs (**A**) and sequences assigned to APases, phosphate transporters (PHT), SPX protein, SIZ1 and PHO1 (**B**). Note that transcripts with RPKM < 3 are presented in Supporting Information Tables S1 and S2. The colour bar indicates the expression levels [represented as log2 (RPKM means)], red colour indicates high expression level while blue indicates low expression level.

### Up-regulated transcripts associated with root-released organic anions

The citric acid and glyoxylate cycles play important roles in synthesis of organic acids in plant tissues. To see if biosynthesis of organic acids could be altered by P deficiency, we mapped the annotated transcripts to genes that involved in citric acid and glyoxylate cycles. Our analysis revealed that 38 up-regulated transcripts identified under P1 treatment represent enzyme-encoding genes putatively involved in the citric acid and glyoxylate cycles (Fig. 5A-B and Table S4). In addition, organic anions were mainly exuded through plasma membrane located transporters. Hence, we further found 10 sequences which were associated with organic anion efflux transporters including the MATE efflux family (transporters that transport a broad range of substrates such as organic anions, plant hormones and secondary metabolites), citrate transporter (CT) and ALMT (aluminium-activated malate transporter), as shown in Fig. 5C and Table S4.

**Figure 5.**
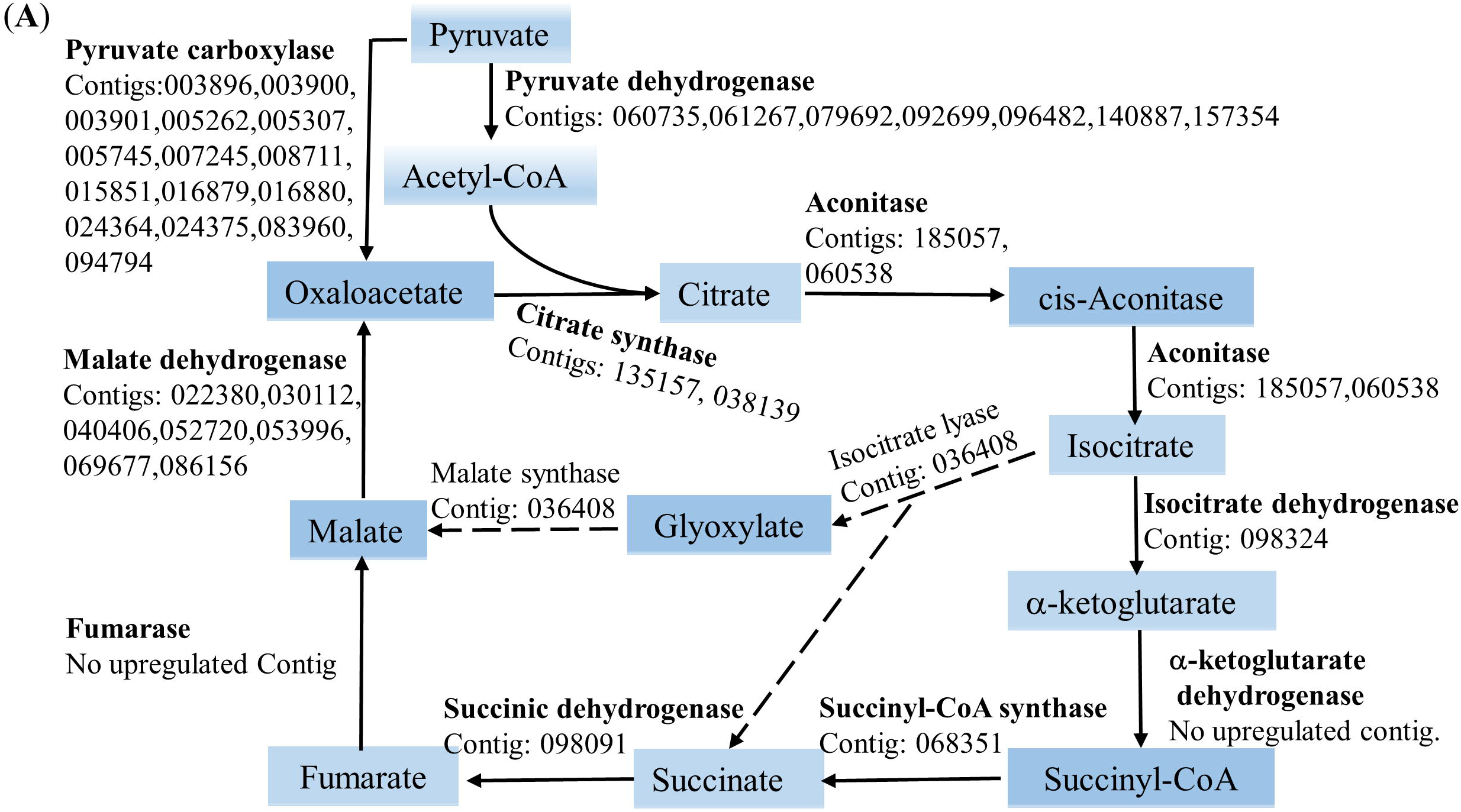
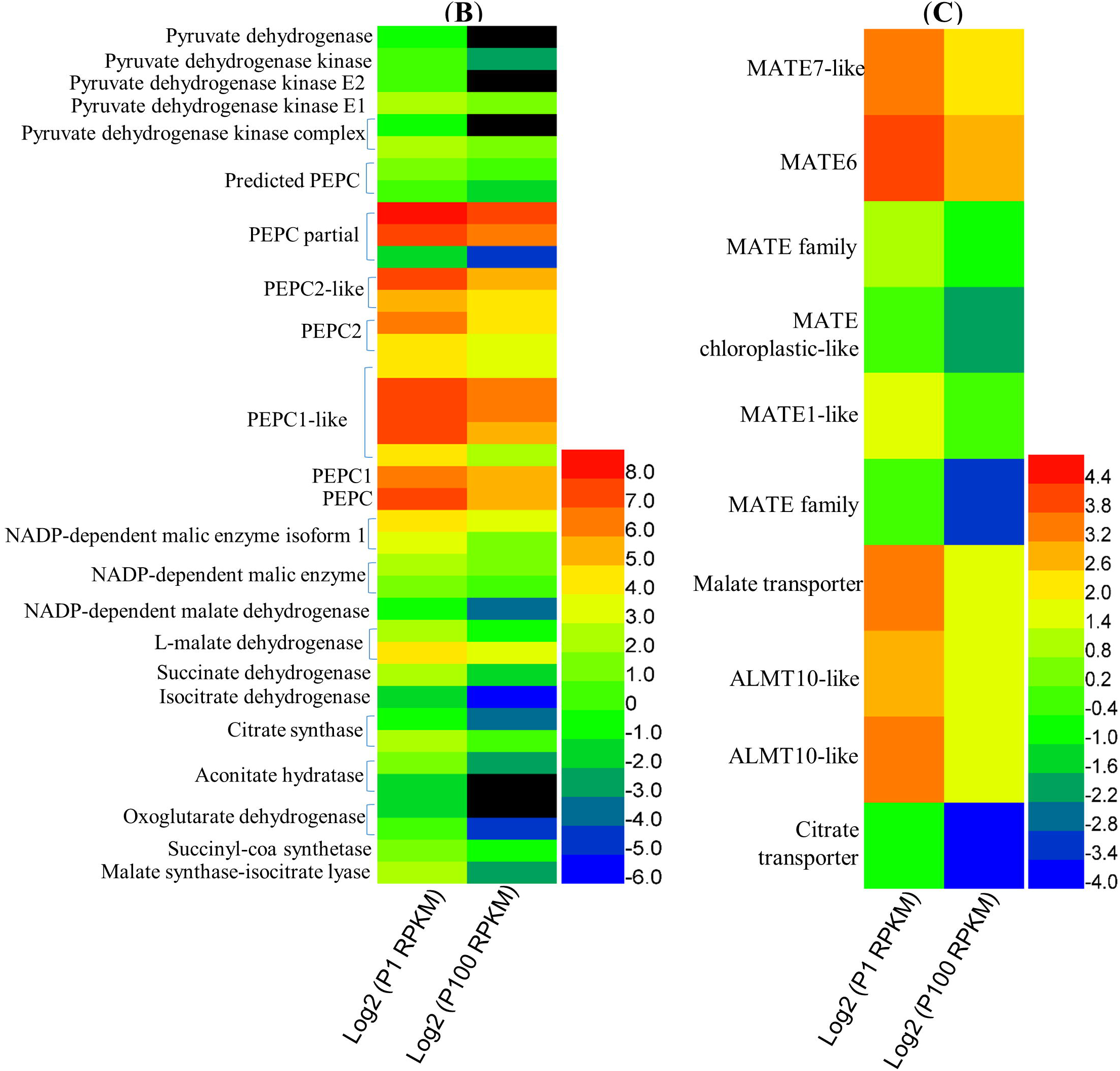
Up-regulated sequences associated with organic anion production and efflux under P deficiency (P1) (**A**) Schematic representation of metabolic pathways including citric acid and glyoxylate cycles related to organic anion production that were up-regulated (*p* < 0.05) under P1 (**B**) and up-regulated organic anion transporters responsive to P deficiency (**C**). The colour bar indicates the expression levels [represented as log2 (RPKM means)], red colour indicates high expression level while blue indicates low expression level and black colour indicates RPKM=0.

### RNA-seq validation by qRT-PCR

To assess whether differential expressed transcripts could be confirmed by an alternate method, 14 transcripts were selected and analysed by qRT-PCR using primers listed in Table S5. Transcripts known to be up-regulated in response to phosphate starvation, i.e. PHO2, PAP3, RNS1 (RNase), PHO1 and SPX, were confirmed by qRT-PCR which showed similar expression patterns to those analyzed by RNA-seq. Additionally, the expression of transcripts involved in root organic anion synthesis such as MSIL (malate synthase-isocitrate lyase), PEPC (phosphoenolpyruvate carboxylase), CS (citrate synthase), LMD (L-malate dehydrogenase), NADP-MD (NADP-dependent malate dehydrogenase) and efflux transporters like ALMT and MATE was also investigated by qRT-PCR. Among 14 transcripts evaluated by qRT-PCR, the trend of changes in 11 (79%) were consistent with the RNA-seq data (Fig. 6).

**Figure 6.**
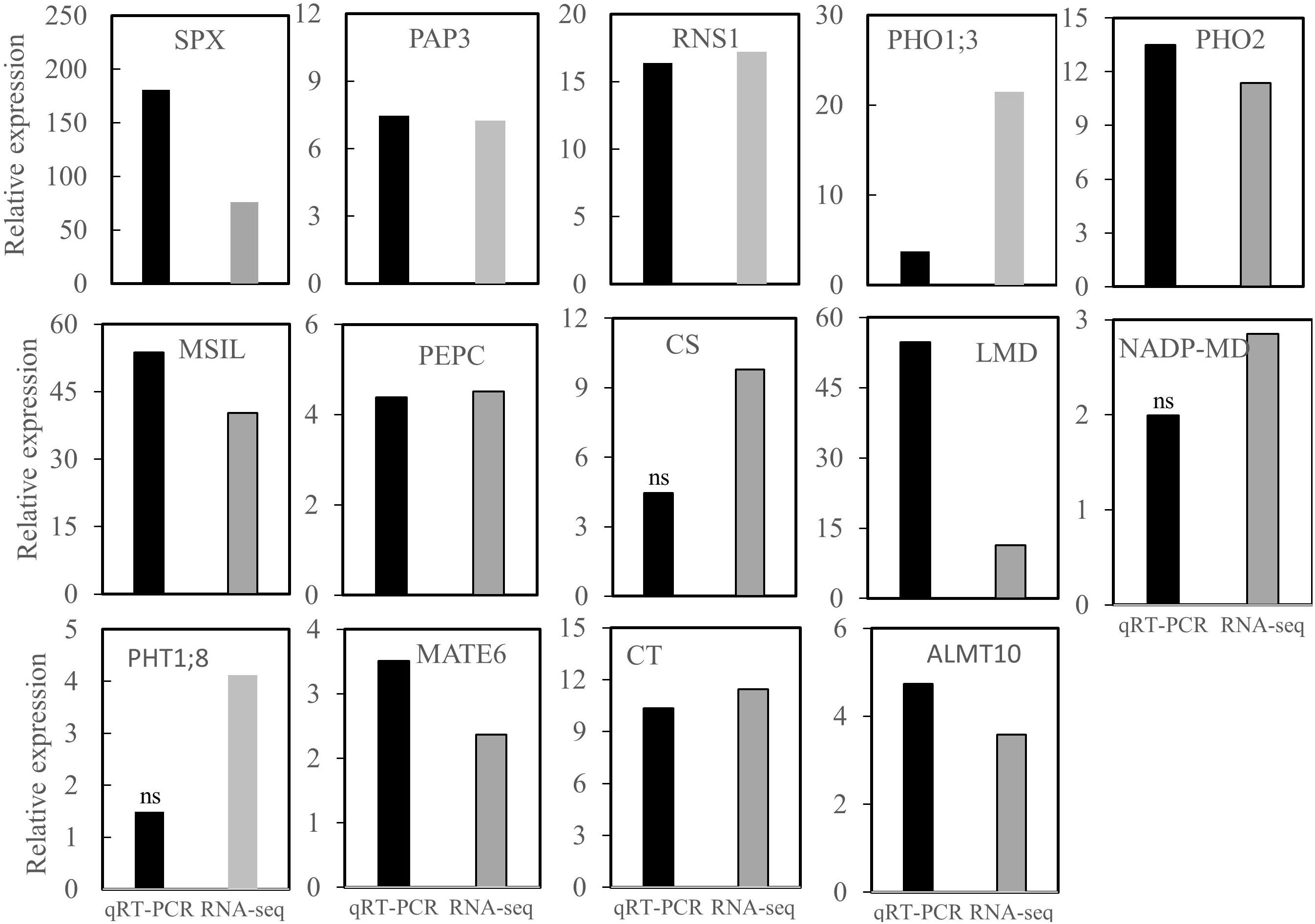
Expression of candidate known genes related to low P stress, and up-regulated transcripts associated with organic anion production and efflux under P1 as determined using RNA-Seq and qRT-PCR. Fourteen genes were selected and analysed using qRT-PCR for both P1 and P100 treatments. Transcript expression levels were normalized using the internal controls β-actin and EF1α (see Methods section). Relative expression values were calculated based on means of four biological replicates (with three technical replicates) under P1 and P100 treatments. Transcripts with statistically insignificant (*p* >0.05) changes in expression compared with P100 roots are denoted as ns. Fold changes based on RPKM values derived from RNA-seq are plotted on the same graph. The transcript IDs for each gene are listed in Supporting Information Table S5.

### Metabolome Analyses

To assess the effects of gene expression in oat roots on overall metabolism, nonbiased metabolite profiling of oat roots was performed using GC-MS. We detected and identified 82 metabolites in oat roots subjected to P1 and P100, as provided in Table S6. Table 4 lists those metabolites that are significantly different (*p*<0.05, *t* test in MeV) between the P1 and P100 treatments as well as the P1/P100 response ratios (based on non-transformed data) and FDR correction (based on Log2 transformed data). The primary metabolites were amino acids, organic acids, polyhydroxy acids, sugars, phosphates, polyols, and *N*-compounds. Most of the metabolites showed a response ratio lower than 1, indicating a decrease in P1 roots; only eight metabolites were increased in P1 roots (Table 4 and Table S6).

**Table 4.** Known metabolites identified by GC-MS in oat roots from P1 and P100 treated plants with *p* < 0.05.

| Class | Metabolite | Response ratio<br>P1/P100 | $p$ value |
| --- | --- | --- | --- |
| **Organic acids** | 2-hydroxy-glutaric acid | 0.10 | 0.0023 |
|  | 2-oxo-glutaric acid | 0.08 | 0.0117 |
|  | Pantothenic acid | 0.69 | 0.0118 |
|  | Pyruvic acid | 0.17 | 0.0101 |
|  | Succinic acid | 0.32 | 0.0063 |
| **Amino acids** | 4-amino-butanoic acid | 0.81 | 0.0209 |
|  | Methionine | 0.25 | 0.0253 |
|  | Valine | 0.35 | 0.0460 |
| **N-compounds** | 5-methylthio-adenosine | 0.22 | 0.0087 |
|  | Putrescine | 0.29 | 0.0063 |
|  | Spermidine | 0.56 | 0.0446 |
| **Phenylpropanoids** | 4-hydroxy-cinnamic acid | 0.75 | 0.0372 |
| **Phosphates** | Ethanolaminephosphate | 0.30 | 0.0404 |
|  | Fructose-6-phosphate | 0.43 | 0.0063 |
|  | Glucose-6-phosphate | 0.19 | **0.0011** |
|  | Glycerophosphoglycerol | 0.27 | 0.0367 |
|  | Mannose-6-phosphate | 0.22 | **0.0011** |
|  | myo-Inositol-phosphate | 0.25 | **0.0016** |
|  | Phosphoric acid | 0.28 | **7.8E-4** |
|  | Phosphoric acid monomethyl ester | 0.41 | 0.0118 |
|  | Glucose-6-phosphate | 0.20 | 0.0034 |
| **Polyhydroxy Acids** | Lyxonic acid | 0.48 | 0.0039 |
|  | Ribonic acid | 0.45 | 0.0087 |
| **Polyols** | Arabitol | 0.56 | 0.0087 |
|  | Inositol, myo- | 0.84 | 0.0157 |
|  | Ribitol | 0.49 | 0.0157 |
| **Sugars** | Sucrose | 0.47 | 0.0357 |
|  | Xylose | 0.65 | 0.0209 |
|  | Glucopyranose | 0.27 | **0.0016** |
|  | Maltose | 0.53 | 0.0207 |

FDR correction with $\alpha < 0.05$ is indicated by bold format of the $p$ value.

PCA analysis of metabolite data using all 82 known metabolites as well as 22 alternative metabolites and 39 mass spectral metabolite tags (MSTs) indicated that PC1 nicely defines the difference between two groups and represents about 35.8% of the variation (Fig.7). However, some overlap in the samples can be seen and high variation within the samples of the same group can be observed, which probably suggested different levels of P deficiency in oat roots and some P100 treated plants might be suffering P deficiency due to rapid depletion of P in the solution. After FDR correction, only five metabolites showed significant differences between P1 and P100 roots: phosphoric acid, mannose-6-phosphate, glucose-6-phosphate, glucopyranose and myo-inositol-phosphate, which indicated that the central metabolism might be stable in oat roots after 10 days of P deficiency. Regarding the organic acids, all identified organic acids showed P1/P100 ratios lower than 1 except for citric and malic acids, which showed P1/P100 ratios of 1.08 and 1.23, respectively (Table 4 and Table S6).

**Figure 7.**
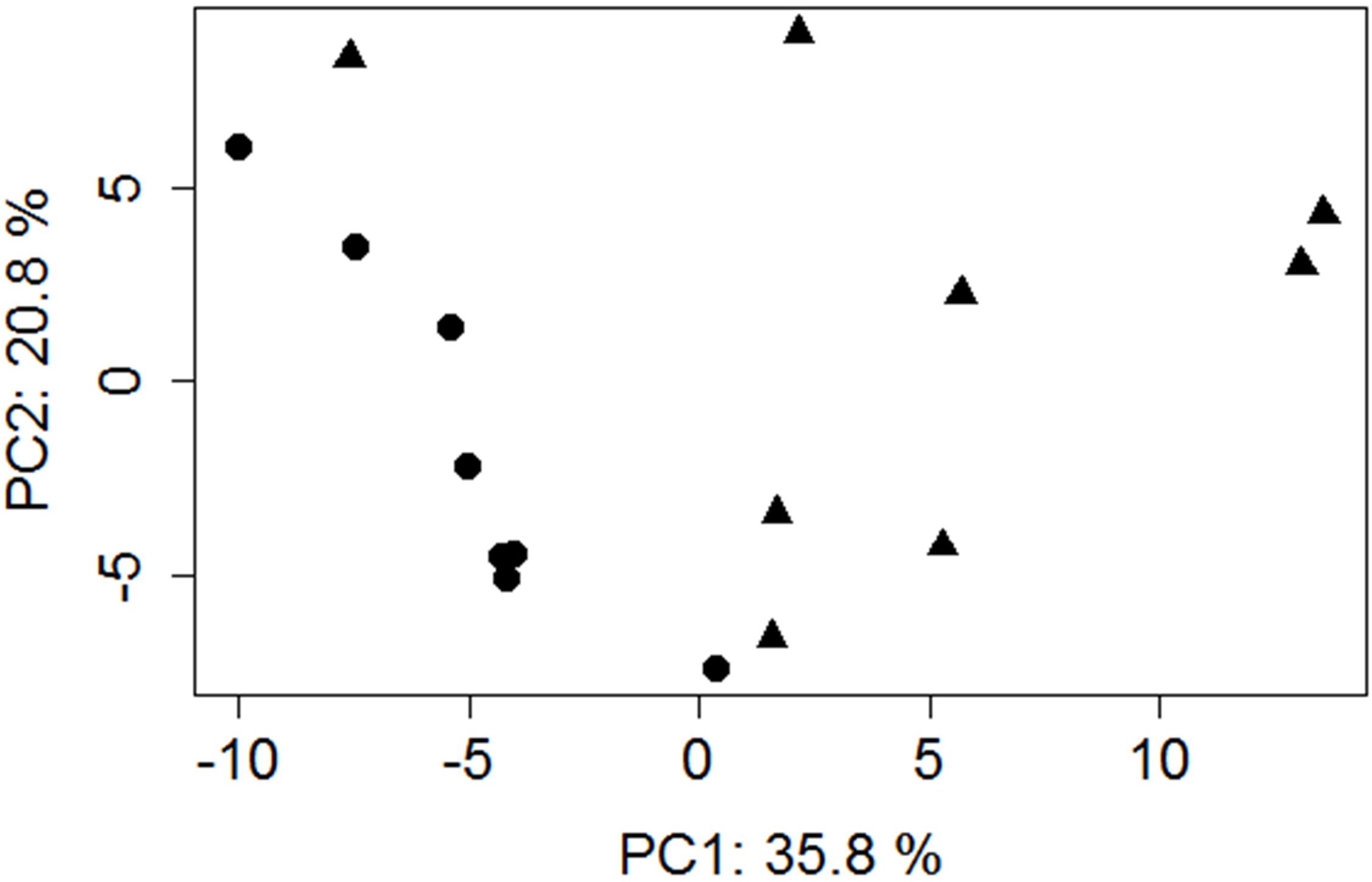
Principal component (PC) scores of metabolic variances in oat roots (n=8 × 2) Oat plants grown in P1(circles) and P100 (triangles) solutions for 10 days were used.

## Discussion

Phosphorus (P) deficiency severely limits plant growth and productivity. This is especially important for sustainable staple cereal crop production in the future. Understanding the molecular mechanisms underlying root and root-secreted organic anion responses to P deficiency in oat, one of the main cereal crops in the world, is of high interest for optimizing future oat production.

Hydroponics as a root environment influences root architecture, in particular root elongation due to reduced mechanical impedance in the absence of solids (Bengough *et al.*, 2001). Previous studies have suggested that root exudation by plants grown in hydroponics is different from root exudation by plants grown in soil (Neumann *et al.*, 2009; Wang *et al.*, 2015 & 2016). Nevertheless, the elimination of other variables such as impact of soil particles and soil microorganisms favour use of hydroponics in root exudation studies (Aulakh *et al.*, 2001; Cheng *et al.*, 2014; Dechassa and Schenk, 2004; Gahoonia *et al.*, 2000; Ligaba *et al.*, 2004). In addition, RNA-seq analysis also benefits from removing the influence of these variables on gene expression, while hydroponics use makes high quality RNA extraction easier.

Oono *et al.* (2011 & 2013) concluded that the highest number of responsive transcripts was observed in roots at 10 days after P deficiency in rice and wheat, while the plants’ morphological and physiological responses to P deficiency become prominent at around 30 days of P starvation (Cheng *et al.*, 2014; Oono *et al.*, 2011; Wang *et al.*, 2015). Hence, we studied transcriptome and metabolome, and root morphology and exudates at different time points (i.e., 10 days for RNA and metabolome samples and two or four weeks for root exudates) after the P1 and P100 treatments. Our gene expression and metabolome profiles represent early to mid-term responses, while the others were mainly long-term P deficiency responses.

Based on our physiological analysis, we did not detect any organic anion exudation after two weeks of P-deficiency in oat. This might be due to: 1) the extracted organic anions were below the detection limit; 2) root released organic anions were affected by the plant developmental stage (Aulakh *et al.*, 2001; Watt and Evans, 1999; Wang *et al.*, 2017). Following four weeks of P-deficiency, under similar growth and sampling conditions, oat root had higher exudation rate of citrate than other species such as canola, rice, cabbage, carrot, barley, soybean and potato (Aulakh *et al.*, 2001; Dechassa and Schenk, 2004; Gahoonia *et al.*, 2000; Ligaba *et al.*, 2004; Liang et al., 2013; Wang *et al.*, 2015), as well as white lupin (Cheng *et al.*, 2014; Watt and Evans, 1999; Wang *et al.*, 2007) which is to our knowledge currently the most efficient species that uses root-secreted citrate to cope with P deficiency (Cheng *et al.*, 2011). Additionally, the high exudation rate of citrate by oat roots under P1 treatment corresponded well with our greenhouse experiment using clay-loam agricultural soils (Wang *et al.*, 2016). Therefore, exudation of citrate appeared to be a late response to P starvation in oat. Given that production and exudation of organic anions is a more carbon-costly process than other pathways (e.g., the production of root hairs and lateral roots) for plants (Lynch, 2007; Whipps, 1990), it might be economical to release organic anions just at a certain stage.

Transcripts encoding PEPC and malate synthase from a glyoxylate-like cycle, which are involved in organic anion production, as well as sequences assigned to citrate and malate efflux transporters were detected in the transcriptome of white lupin cluster roots under low P stress (O’Rourke *et al.*, 2013). By contrast, such transcripts were not reported in wheat, rice, *Arabidopsis* or potato studies (Hammond *et al.*, 2011; Oono *et al.*, 2011; Oono *et al.*, 2013; Misson *et al.*, 2005), probably due to these plants exhibiting a low exudation rate of organic anions under P deficiency (Aulakh *et al.*, 2001; Neumann and Romheld, 1999; Narang *et al.*, 2000; Wang *et al.*, 2015). In P starved oat roots, we identified 38 up-regulated transcripts encoding almost all enzymes associated with the citric acid and glyoxylate cycles except for fumarase and α-ketoglutarate dehydrogenase. Moreover, a transcript annotated as malate synthase-isocitrate lyase (MSIL) was highly expressed (> 40-fold) under the P1 compared with the P100 treatment, suggesting an important role of the glyoxylate cycle in organic anion production in oat. This gene is of interest and could be used to improve P uptake in other species using genetic engineering. Exudation of organic anions may also lead to alteration of gene expression of enzymes involved in organic acid metabolism, but this is unlikely to be the case in the current study since no organic anion exudation was detected yet when we sampled RNA.

In white lupin, enhanced levels of citrate were observed in roots (2.2 fold) and cluster roots (7.6 fold) after 22 days P deficiency, whereas after 14 days P deficiency the changes were 1.4 and 3.5 fold, respectively (Müller *et al.*, 2015), suggesting that changes in the metabolome mainly occurred after long-term P deficiency. Our oat root metabolome analysis indicated that most organic acids showed a general reduction after 10 days of P deficiency, which corresponded well with common bean roots after 21 days of low-P treatment (Hernández *et al.*, 2007), while slight (but not significant) increases in citric and malic acids were detected in our study. Previous studies have suggested that biosynthesis and exudation of organic anions has been associated with enhanced expression of genes encoding PEPC, malate dehydrogenase, citrate synthase and transporters like ALMT and MATE (de la Fuente *et al.*, 1997; Delhaize *et al.*, 2009; Johnson *et al.*, 1994; Koyama *et al.*, 2000; Watt and Evans, 1999; Wang *et al.*, 2013). However, interpretation of links between gene expression and organic acids biosynthesis and exudation should be done with caution, because enhanced gene expression does not necessarily result in enhanced enzyme levels (and enzyme activities). Also, other cellular conditions caused by P deficiency can affect endogenous enzyme function (Ryan *et al.*, 2001). Additionally, although a number of studies have shown associations between organic anion efflux and internal concentrations (Hoffland *et al.*, 1989; Neumann and Romheld, 1999), internal concentrations of organic anions are unlikely to directly regulate organic anions efflux in P-deficient plants (Keerthisinghe *et al.*, 1998; Ryan *et al.*, 2001; Watt and Evans, 1999). Rather, transporters are likely to be the most important regulators of organic anion exudation (Ryan *et al.*, 2001). We identified ten up-regulated transcripts encoding MATE and ALMT family members, and other citrate and malate transporters. While higher expression of transporter-encoding genes may increase the number of transporters per cell, the expression of these transcripts was not high (0.66–16.66 RPKM), and enhanced transcript accumulation cannot be assumed to equal increased protein abundance. Furthermore, efflux is determined by both abundance and activity, with regulation of the latter still largely unknown.

Among the known genes expressed in P deficiency, a highly conserved PHR1-IPS1-miRNA399-PHO2 signalling cascade has been elucidated in Arabidopsis and rice (Lin *et al.*, 2009; Oono *et al.*, 2013). PHR1 (PHR2 in rice) is a MYB-type transcription factor, acting as a key factor in regulating downstream P deficiency responsive gene expression. Both AtPHR1 and OsPHR2 were not very responsive to P deficiency, but their overexpression activated the expression of a number of P-starvation induced genes even under P sufficient conditions (Rubio *et al.*, 2001; Zhou *et al.*, 2008). We did not identify any transcript annotated as PHR1 or PHR2 in our *dn*ORT database. We detected up-regulated SPX, PHO2, RNS1 and SIZ1 in oat. SPX may inhibit the expression of PHR1, and SIZ1 facilitates sumoylation of PHR1 and thereby regulates the post-translational modification of PHR1 (Chiou & Lin, 2011; Wu *et al.*, 2013), which likely explains why we could not detect differentially expressed PHR1 in oat and suggests that the PHR1-IPS1-miRNA399-PHO2 signalling cascade is likely also conserved in oat.

In our study, we also detected many CCAAT-box binding transcription factors, including Nuclear Factor (NF) Y subunits NF-YA, NF-YB and NF-YC, which respond to P deficiency in oat. CCAAT-box transcription factors, in particularly NFYA-B1, play essential roles in root development and P uptake in wheat (Qu *et al.*, 2015). Our previous study also found that root morphology, rhizosphere bacteria and root-colonizing mycorrhizal fungi were involved in the response to low P availability in oat (Wang *et al.*, 2016). The current study identified about 30 up-regulated transcripts associated with auxin responses, which might regulate root morphology; more than 60 transcripts involved in disease response and 9 involved in responses to fungal infection. Additionally, 24 up-regulated transcripts under P deficiency found in the present study had been reported previously in rice and wheat (Oono *et al.* 2011 & 2013), suggesting that these genes are valuable indicators of P deficiency in cereal crops. Another 25 unique transcripts in oat that were up-regulated more than two-fold under P deficiency were identified. These will be studied further to investigate their roles related to P uptake in oat, in order to facilitate future improvements in oat production.

In summary, our current study provides new insights into the molecular mechanisms involved in root responses to P deficiency and the release of organic anions by P-starved oat roots. The novel information generated in the present study enriches our understanding of oat adaptation to low P availability and contributes to future sustainable oat production.

## Acknowledgements

This study was supported by the strategic institute program (SIS) on “Opportunities for sustainable use of phosphorus in food production” at the Norwegian Institute of Bioeconomy Research (NIBIO). The authors thank Marit Almvik and Monica Skogen for their kind help with LC-MS/MS and RNA sample preparation, respectively. We are grateful to Prof. Nicholas Clarke for linguistic correction and to the Norwegian Sequencing Centre, Oslo, Norway, for library preparation and sequencing. The Norwegian Sequencing Centre, a national technology platform hosted by the University of Oslo and supported by the “Functional Genomics” and “Infrastructure” programs of the Research Council of Norway and the South-eastern Regional Health Authorities, provided the sequencing service. We thank Ines Fehrle and Joachim Kopka (both from Max Planck Institute of Molecular Plant Physiology, DE) for their excellent technical assistance and support in analysis of oat metabolites.

## Author Contributions

Y.W. contributed to the experimental design, sample preparation, plant biomass and root morphology measurements, data analyses and manuscript writing; J.L.C. contributed to the experimental design and conceived the study; E.L. did the sequence trimming, *de novo* assembly, annotation, and helped with the data analyses; T.A-M and A.E conducted metabolite measurements; L.P. helped with the qRT-PCR experiment. All authors revised and approved the final manuscript.

## Supplementary Data

The following supplementary data is available for this article:

**Fig. S1** Data distribution of Blastx hits of *dn*ORT sequences.

**Fig. S2** The Blastx top-hit species distribution of *dn*ORT sequences.

**Fig. S3** Functional gene onthology (GO) classification of *dn*ORT sequences.

**Fig. S4** Putative functions (with InterProScan) distribution of *dn*ORT sequences.

**Table S1.** Up-regulated transcription factors under P deficiency.

**Table S2.** Up-regulated transcripts predicted to be acid phosphatases (APases), phosphate transporters and other known genes related to P deficiency.

**Table S3.** Up-regulated transcripts associated with auxin responses, disease responses and responses to fungal infection under P deficiency.

**Table S4.** Up-regulated transcripts associated with organic anion production and efflux under P deficiency.

**Table S5.** Primers used in the present study.

**Table S6.** GC-MS metabolite profiles.

